# Distribution of duck-origin parvovirus over time in Cherry Valley ducks in vivo and histopathological investigation

**DOI:** 10.1101/420067

**Authors:** Bing Chen, Qihui Luo, Jing Xu, Chao Huang, Wentao Liu, An-chun Cheng, Zhengli Chen

## Abstract

In 2015, we successfully isolated a strain of duck-origin parvovirus from Cherry Valley ducks, which we named QH-L01. In this study, duck-origin parvovirus in Cherry Valley ducks was quantified and localized by quantitative real-time PCR (qPCR) and immunohistochemistry (IHC), and pathological damage to the tissues and organs was observed by hematoxylin-eosin staining (HE staining). qPCR showed that the viral load was higher in the spleen, brain, lung, cecum, ileum, and duodenum over time. The results from IHC experiments showed positive reactions in hepatocytes, epithelium of the lung atrium, myocardial cells, goblet cells of the intestine, and brain cells. Primary histological examination revealed pulmonary lobule depletion and dilation in the lung as well as necrosis and erosion of the villus tips in the duodenum, ileum and cecum. This study is the first demonstration that duck-origin parvovirus can be transmitted from the spleen to the brain and lung, resulting in proliferation and dissemination of the virus to the cecum, ileum, duodenum and other tissues through the blood. The lung, duodenum, ileum and cecum may thus represent the main target tissues and organs for duck-origin parvovirus.

## 1. Introduction

In March 2015, a newly epizootic disease outbreak was observed in Cherry Valley ducks in China that caused beak atrophy and dwarfism syndrome (BADS) [1]. According to relevant reports, the etiological agent of BADS was a novel duck-origin parvovirus similar to Muscovy duck parvovirus (MDPV) and goose parvovirus (GPV) and belonged to the genus *Dependoparvovirus* of the family Parvoviridae [2-3]. The morbidity in Cherry Valley ducks with typical symptoms of BADS has been reported as 15% to 25% and even up to 50% [4]. The diseased Cherry Valley ducks suffer from obvious growth retardation with serious weight loss and beak atrophy, and the other clinical symptoms included swollen tongue, watery diarrhea, degenerative loss of skeletal muscle, and even death [5]. Over the last two years, this virus has caused considerable economic loss for the intensive duck production systems of China.

Representative BADS isolates containing SDLC01 [1], SBDSV-M15 [6], AH-D15 [7], SD [8], and GPV-QH15 [9] have been reported in mainland China since March 2015. In April 2016, the QH-L01 virus, which causes characteristic clinical symptoms of BADS, was successfully isolated from parvovirus antibody-free embryonated Cherry Valley duck eggs and was thoroughly described in another paper [10]. Initially, the causative agent of BADS was considered a variant of GPV [11]; those BADS isolates are closely related to GPV on the basis of the reported phylogenetic analysis [4]. Nevertheless, compared to acute GPV, the BADS isolates obviously differ in the resulting clinical symptoms and mortality and morbidity rates. GPV usually causes ascites, enteritis, myocarditis, and hepatitis with high mortality and morbidity in Muscovy ducklings and goslings and is widely known as Derzsy’s disease or gosling plague in China [12-13].

There are many reports that have analyzed pathogenesis by quantitative real-time PCR (qPCR) and damage to organs and tissues by HE staining for other viruses, such as GPV [14] and enterovirus 71 [15]. However, currently no relevant data have been reported about the distribution and lesions of duck-origin parvovirus to tissues and organs following experimental infection. In addition, the dynamic distribution and nature of the damage to organs and tissues caused by duck-origin parvovirus is unknown. Therefore, the proliferation and distribution of duck-origin parvovirus and the related histopathological changes in vivo requires further study. qPCR is one of the most effective methods for quantitative studies of viral load in recent years [16-17]. IHC is one of the common methods used for antigen localization in cells [18]. HE staining can be used to evaluate the pathological lesions in cells [19-20]. Therefore, using the three described techniques, we studied the dynamic distribution of duck-origin parvovirus and pathological damage to organs and tissues in Cherry Valley ducks after experimental infection to provide guidance for the early prevention of this disease [21].

## 2. Materials and Methods

### 2.1 Virus isolation

The strains of duck-origin parvovirus used in this study were isolated from a Cherry Valley duckling presenting symptoms of BADS in the Sichuan Province of China in April 2016. The QH-L01 virus was successfully isolated from a 9-day-old Cherry Valley duck embryo through the allantoic cavity, and ELD_50_ was 10^-6.54^.

### 2.2 Ethics Statement

Cherry Valley ducks (2 days old) for experiments were obtained from the breeding facility of the Institute of Poultry Sciences at Sichuan Agricultural University in China. The animals were bred and maintained in an accredited facility at the same institution, the experiments conformed to the principles outlined by the Animal Care and Use Committee of Sichuan Agricultural University, and experiments were performed in accordance with the “Guidelines for Experimental Animals” of the Ministry of Science and Technology (Beijing, China). The Animal Care and Use Committee of Sichuan Agricultural University approved our experiments (SYXK(Chuan)2014-187). The ducks were euthanized by CO2 for 10 minutes before sample collection. The experimental ducks monitored for adverse clinical signs were made three times daily. Once the ducks were breathing shallow and the apathetic, it was judged to be dying and has been put to sleep. Ducks that died in the course of this study (i.e. the 20% of individuals in the experimental group) humanely euthanised at the onset of humane endpoint clinical signs. All research staff performed experiments according to the approved protocol. All efforts were made to minimize suffering. A neutralization test was conducted to confirm that the Cherry Valley ducks did not have antibodies against duck-origin parvovirus before the experimental infections.

### 2.3 Infection of Cherry Valley ducks with duck-origin parvovirus

Seventy Cherry Valley ducks were randomly divided into 2 groups. Briefly, 45 Cherry Valley ducks were intramuscularly inoculated with 0.2 mL of 10^6.54^ ELD_50_ duck-origin parvovirus strain. Another 25 Cherry Valley ducks were treated with an equal volume of physiologic saline and used as controls. Three ducks from the infected group and two ducks from the control group were euthanized at each time point. The blood, heart, liver, spleen, lung, kidney, pancreas, proventriculus, gizzard, tongue, thymus, esophagus, trachea, brain, duodenum, jejunum, ileum, cecum, and rectum were then collected for viral load detection and histopathological investigation at different postinoculation (PI) time points: 0.5, 1, 2, 4, 8, 12, 24, 48, 72, 144 and 216 h. The tissues were divided into two approximately equal portions. One portion was frozen at −80 centigrade, weighed, and homogenized using an Omni PCR Tissue Homogenizer (Omni). The other portion was fixed in 4% neutral formaldehyde. For the assays, tissue samples were homogenized in 1 mL of PBS (pH=7.4). DNA was extracted from the tissue samples by using a TIANNamp Genomic DNA Kit (Tiangen Biotech (Beijing) Co., Ltd.). This assay was used to quantify the viral load. Assays for all the samples were repeated three times. The viral concentrations were expressed as the mean log10 of virus genome copy numbers per g or 1 mL of the tested tissue or blood, respectively. In addition, the fixed tissues were submitted for routine paraffin sectioning for IHC and HE staining by LEICA RM2135, and microscopic examination was used to observe virus location and pathological form.

### 2.4 Histopathological and Immunohistochemical Analyses

Various tissue samples from the organs and tissues of infected Cherry Valley ducks, including the heart, liver, spleen, lung, kidney, pancreas, proventriculus, gizzard, tongue, thymus, esophagus, trachea, brain, duodenum, jejunum, ileum, cecum, and rectum, were fixed in 4% neutral paraformaldehyde, dehydrated using an ethanol gradient, and embedded in paraffin before obtaining 5 μm sections for further HE staining. Histopathological analysis of the tissue sections from each organ was performed under a light microscope.

For immunohistochemical analysis, the samples were paraffin embedded, sectioned (5 μm), and placed on poly-L-lysine-coated glass slides, and the endogenous peroxidase activity of the tissues was inhibited by treatment with hydrogen peroxide. The samples were preincubated in 10% normal goat serum in PBS containing 1% bovine serum albumin (BSA) and 0.3% Triton X-100 for one hour at room temperature to block nonspecific binding activity. Then, the samples were incubated overnight (4 centigrade) with anti-goose parvovirus VP3 antibody (1:200) (Bioss, Beijing, China). Then, the sections were incubated with goat anti-rabbit IgG antibody (1:100) (Boster, Wu Han, China) at room temperature for one hour. The antigen-antibody reaction for immunohistochemical staining was performed using a standard avidin-biotin-peroxidase complex technique (IHC kit, Boster, China), and PBS was used as a negative control.

### 2.5 Viral DNA Extraction and qPCR Assay

Viral DNA was extracted from 0.05 g of fresh tissues and 200 μL of blood from the experimental animals (for the spleen only 0.01 g) and controls using a TIANNamp Genomic DNA Kit according to the manufacturer’s protocol (Tiangen Biotech (Beijing) Co., Ltd.). qPCR assays were performed using the Applied Biosystems^®^ 7500 FAST Real-Time PCR System and SYBR Green Master Mix. The reactions, data acquisition, and analyses were performed using Bio-Rad CFX96^TM^ manager software. The qPCR assay was performed in a 25.0 μL reaction mixture that contained 12.5 μL SYBR Green Master Mix, 1.0 μL forward primer (F: CACCACCACAGGTCTTCATCAA), 1.0 μL reverse primer (R: GCTGGTGAACTGAATTTCTGGAT), and 1.0 μL DNA template according to the manufacturer’s instructions, and ddH2O was added to bring the final volume to 25.0 μL. The qPCR amplicon size was 160 bp. The tests were performed using 0.2 mL PCR tubes (ABgene, UK). A standard reference curve was obtained by measurements of serially diluted recombinant plasmid DNA (pVP1) by ligating the PCR product into the pGM-Simple-T Fast Vector (Tiangen Corp, Beijing, China) and transforming *E. coli* DH5α competent cells with the resulting vector [22]. On the basis of the molecular weight, we calculated the plasmid DNA VP1 copy number using the equations described by Ke [23].

### 2.6 Statistical Analysis

Data are expressed as the average of three samples in some experiments. Typically, five slices were selected for each tissue/organ for immunohistochemistry and pathological observation.

### 2.7 Data availability statement

The data sets generated during and/or analyzed during the current study are available from the corresponding author on reasonable request.

## 3. Results

### 3.1 Clinical symptoms of duck-origin parvovirus infection

In this study, the duck-origin parvovirus virus isolate QH-L01 was inoculated into 2-day-old healthy Cherry Valley ducks. The behavior of some of the Cherry Valley ducks presented as depression, mental fatigue, dyspnea, diarrhea and even the initial stages of death after 2 days. The infected ducks had a low mortality rate (20%); 9 ducks died from 2 to 6 days postinoculation, with punctate hemorrhage observed in the beak (Table 1). The necropsies of the dead ducks showed systemic hemorrhage, including lung hemorrhaging and duodenum hemorrhaging with effusion. In contrast, no obvious changes were observed in the control group.

**Table 1.**
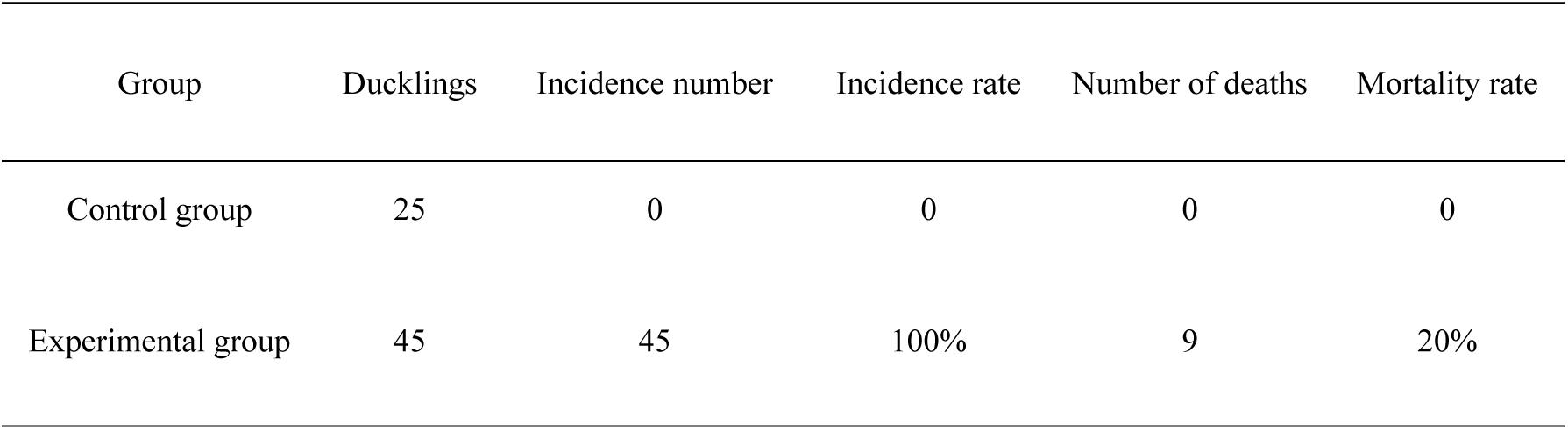
The number of dead ducks

### 3.2 Dynamic distribution of duck-origin parvovirus in vivo

There was a linear correlation between the Ct value and the logarithm of the plasmid copy number (R^2^=0.996), and the copies of virus DNA were calculated according to the standard curve equation: Ct=-3.480×lg [virus copies]+39.959 (R^2^=0.996). According to the Ct value of each sample in the standard curve, the initial viral DNA copy number of the sample was calculated by using the established qPCR. The qPCR data from all the organs and tissues are shown in Table 2. Select organs containing the highest copy number of virus at each time point were selected for further study (Fig 1). The results revealed that the virus was first detected in the spleen and blood at 1 h PI, and the virus copy numbers were 10^2.86^ copies/g and 10^2.62^ copies/mL. At 2-4 h PI, the heart, liver, lung, kidney, pancreas, trachea, tongue, esophagus, gizzard and brain were positive for virus. At 8 h PI, duck-origin parvovirus can be detected in all tissues and organs. It is worth noting that at different time points of infection, the tissue or organ with the maximum copy of virus is different. At 4 h PI, the tissue or organ with maximum copies of virus is the spleen, with 10^6.00^ copies/g. At 8 h PI, it is brain, with 10^6.11^ copies/g, and then lung (12 h), cecum (24 h), ileum (48 h), duodenum (72 h-216 h), with 10^8.28^ copies/g, 10^9.23^ copies/g, 10^7.1^ copies/g, 10^8.54^ copies/g, 10^9.83^ copies/g, and 10^9.01^ copies/g, respectively. Throughout infection, QH-L01 DNA was detected in the blood. Interestingly, from 4 h-216 h PI, in the main tissues and organs, including the heart, liver, spleen, lung, kidney, duodenum, jejunum, ileum and cecum, higher viral loads of duck-origin parvovirus were seen than in the blood, and the virus DNA copies were stable in the blood at approximately 10^4.5^ copies/mL. The general trend of QH-L01 DNA copies increased at 24 h-48 h and decreased at 48 h-144 h, at which time the viral proliferation in the Cherry Valley ducks induced immune responses to eliminate virus particles except for in intestinal tissues at 48 h PI. The control group was negative for viral detection.

### 3.3 Histopathological changes caused by duck-origin parvovirus infection in ducks

According to the qPCR results, we chose two time points to analyze tissues and organs histopathological changes, and performed paraffin section with HE staining. One of the time points was the maximum virus copy number, and we performed microscopic examination demonstrating that the spleen, brain, lung, cecum, ileum and duodenum were damaged at the different time points. The spleen, as one of the immune organs, showed lymphocyte infiltration at 48 h PI, which indicated that the immune response began to strengthen (Fig 2 A4, arrow). However, there was no obvious pathological injury at 4 h PI (Fig 2 A2). Under the meninges, we observed inflammatory cell infiltration from 48 h PI (Fig 2 B4), but not in the brain at 8 h PI (Fig 2 B2). In the pulmonary lobule, we observed serious compensatory dilation, and the number of lung capillaries decreased markedly at 12 h PI (Fig 2 C2). Furthermore, the lung presented various degrees of hemorrhage in the experimental group at 72 h PI (Fig 2 C4). The top of the cecum, ileum and duodenal villi showed gradual necrosis or erosion in the experimental group at 24 h, 48 h and 72 h (Fig 2 D2, E2 and F2). In addition, compared with the control group, the lymphocyte numbers in the cecum increased from 24 h - 216 h PI (Fig 2 D4), and many inflammatory cell infiltrates were observed in the tissue of the ileum at 72 h PI (Fig 2 E4). There were no obvious pathological changes in the heart, liver and other organs/tissues of virus-infected Cherry Valley ducks (Fig 3). No obvious pathological injury was observed in the control group.

### 3.4 Viral Antigen Expression in Some Organs of duck-origin parvovirus-infected Cherry Valley ducks

To further investigate the relationship between duck-origin parvovirus load in tissues and its replication, the expression of duck-origin parvovirus antigen in the associated organs of infected Cherry Valley ducks was investigated. According to the results of qPCR and HE staining, we choose the spleen, brain, lung, cecum, ileum and duodenum as tissues to investigate by IHC. There was high expression of the duck-origin parvovirus antigen in the spleen, brain, lung, cecum, ileum and duodenum. In the spleen, the white pulp and red pulp both had positive expression (Fig 4 A2). In the brain, there was positive expression in brain cells (Fig 4 B2). In the lung, there was positive expression of duck-origin parvovirus antigen in the epithelium of the lung atrium (Fig 4 C2). Goblet cells of the villus and the intestinal crypt of the cecum, ileum and duodenum all had positive expression (Fig 4 D2, E2, F2). To further explore whether the virus had also entered the cells of other organs, the liver, heart, kidney and jejunum were analyzed by immunohistochemistry. The results showed that the positive antigen labeling was detected in the liver, heart, kidney and jejunum (Fig 5). No obvious antigen signals were observed in the control group.

**Table 2.**
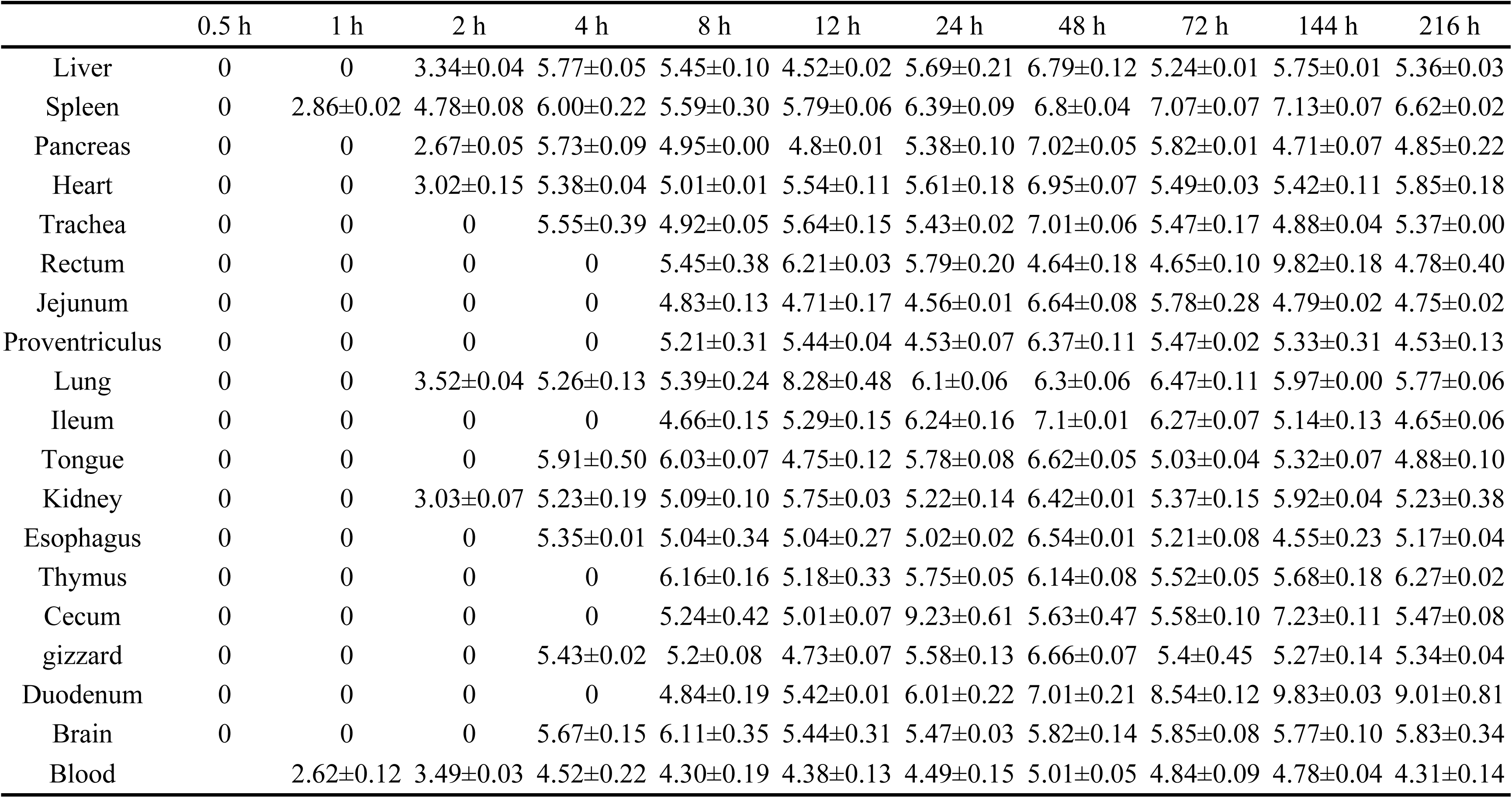
The distribution and quantity of duck-origin parvovirus at different time points within the different segments of the tissue samples

|  | 0.5 h | 1 h | 2 h | 4 h | 8 h | 12 h | 24 h | 48 h | 72 h | 144 h | 216 h |
| --- | --- | --- | --- | --- | --- | --- | --- | --- | --- | --- | --- |
| Liver | 0 | 0 | 3.34±0.04 | 5.77±0.05 | 5.45±0.10 | 4.52±0.02 | 5.69±0.21 | 6.79±0.12 | 5.24±0.01 | 5.75±0.01 | 5.36±0.03 |
| Spleen | 0 | 2.86±0.02 | 4.78±0.08 | 6.00±0.22 | 5.59±0.30 | 5.79±0.06 | 6.39±0.09 | 6.8±0.04 | 7.07±0.07 | 7.13±0.07 | 6.62±0.02 |
| Pancreas | 0 | 0 | 2.67±0.05 | 5.73±0.09 | 4.95±0.00 | 4.8±0.01 | 5.38±0.10 | 7.02±0.05 | 5.82±0.01 | 4.71±0.07 | 4.85±0.22 |
| Heart | 0 | 0 | 3.02±0.15 | 5.38±0.04 | 5.01±0.01 | 5.54±0.11 | 5.61±0.18 | 6.95±0.07 | 5.49±0.03 | 5.42±0.11 | 5.85±0.18 |
| Trachea | 0 | 0 | 0 | 5.55±0.39 | 4.92±0.05 | 5.64±0.15 | 5.43±0.02 | 7.01±0.06 | 5.47±0.17 | 4.88±0.04 | 5.37±0.00 |
| Rectum | 0 | 0 | 0 | 0 | 5.45±0.38 | 6.21±0.03 | 5.79±0.20 | 4.64±0.18 | 4.65±0.10 | 9.82±0.18 | 4.78±0.40 |
| Jejunum | 0 | 0 | 0 | 0 | 4.83±0.13 | 4.71±0.17 | 4.56±0.01 | 6.64±0.08 | 5.78±0.28 | 4.79±0.02 | 4.75±0.02 |
| Proventriculus | 0 | 0 | 0 | 0 | 5.21±0.31 | 5.44±0.04 | 4.53±0.07 | 6.37±0.11 | 5.47±0.02 | 5.33±0.31 | 4.53±0.13 |
| Lung | 0 | 0 | 3.52±0.04 | 5.26±0.13 | 5.39±0.24 | 8.28±0.48 | 6.1±0.06 | 6.3±0.06 | 6.47±0.11 | 5.97±0.00 | 5.77±0.06 |
| Ileum | 0 | 0 | 0 | 0 | 4.66±0.15 | 5.29±0.15 | 6.24±0.16 | 7.1±0.01 | 6.27±0.07 | 5.14±0.13 | 4.65±0.06 |
| Tongue | 0 | 0 | 0 | 5.91±0.50 | 6.03±0.07 | 4.75±0.12 | 5.78±0.08 | 6.62±0.05 | 5.03±0.04 | 5.32±0.07 | 4.88±0.10 |
| Kidney | 0 | 0 | 3.03±0.07 | 5.23±0.19 | 5.09±0.10 | 5.75±0.03 | 5.22±0.14 | 6.42±0.01 | 5.37±0.15 | 5.92±0.04 | 5.23±0.38 |
| Esophagus | 0 | 0 | 0 | 5.35±0.01 | 5.04±0.34 | 5.04±0.27 | 5.02±0.02 | 6.54±0.01 | 5.21±0.08 | 4.55±0.23 | 5.17±0.04 |
| Thymus | 0 | 0 | 0 | 0 | 6.16±0.16 | 5.18±0.33 | 5.75±0.05 | 6.14±0.08 | 5.52±0.05 | 5.68±0.18 | 6.27±0.02 |
| Cecum | 0 | 0 | 0 | 0 | 5.24±0.42 | 5.01±0.07 | 9.23±0.61 | 5.63±0.47 | 5.58±0.10 | 7.23±0.11 | 5.47±0.08 |
| gizzard | 0 | 0 | 0 | 5.43±0.02 | 5.2±0.08 | 4.73±0.07 | 5.58±0.13 | 6.66±0.07 | 5.4±0.45 | 5.27±0.14 | 5.34±0.04 |
| Duodenum | 0 | 0 | 0 | 0 | 4.84±0.19 | 5.42±0.01 | 6.01±0.22 | 7.01±0.21 | 8.54±0.12 | 9.83±0.03 | 9.01±0.81 |
| Brain | 0 | 0 | 0 | 5.67±0.15 | 6.11±0.35 | 5.44±0.31 | 5.47±0.03 | 5.82±0.14 | 5.85±0.08 | 5.77±0.10 | 5.83±0.34 |
| Blood |  | 2.62±0.12 | 3.49±0.03 | 4.52±0.22 | 4.30±0.19 | 4.38±0.13 | 4.49±0.15 | 5.01±0.05 | 4.84±0.09 | 4.78±0.04 | 4.31±0.14 |

**Fig 1.**
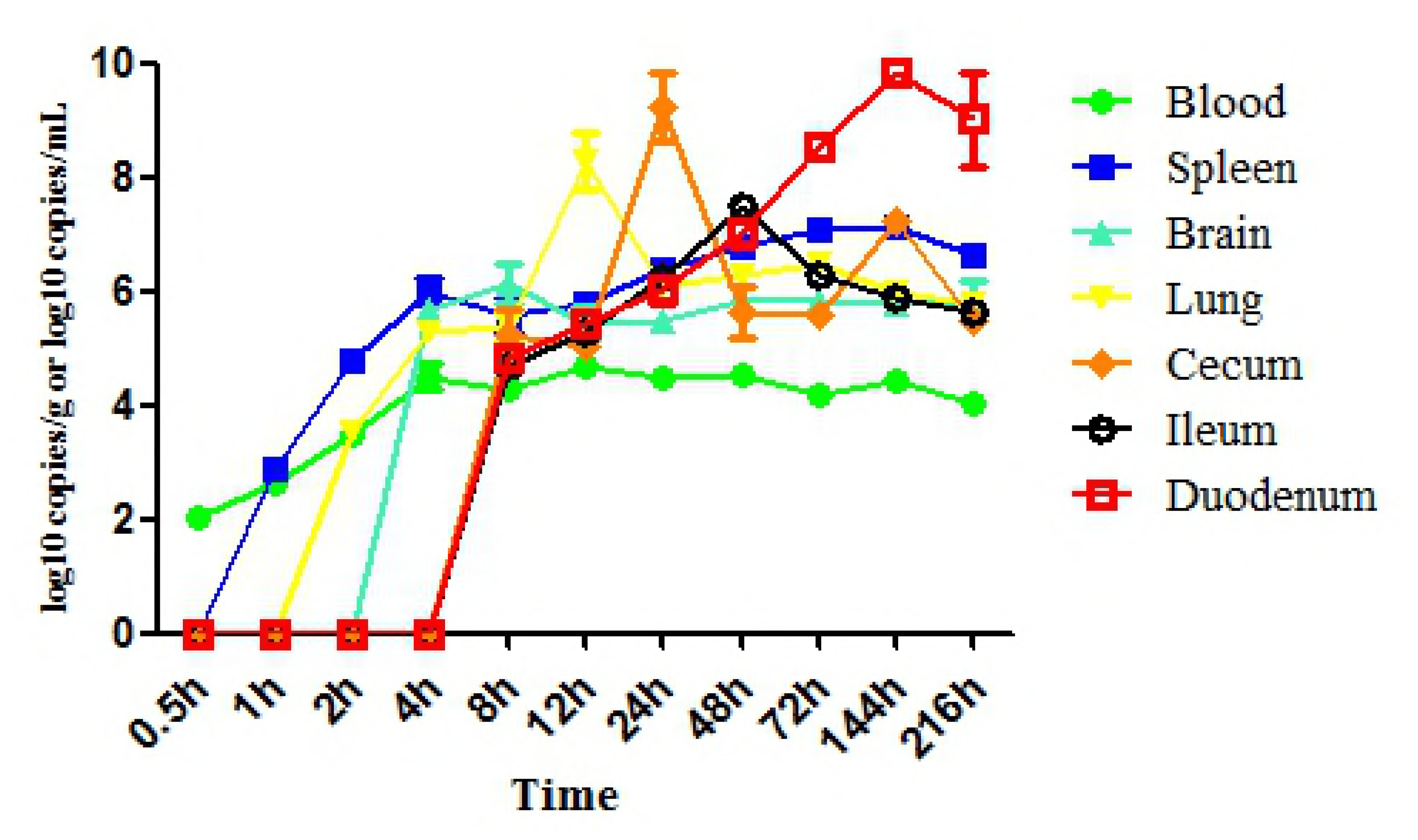
Viral load in tissue samples from Cherry Valley ducks infected with QH-L01. The values are the means for three replicates at each time point

**Fig 2.**
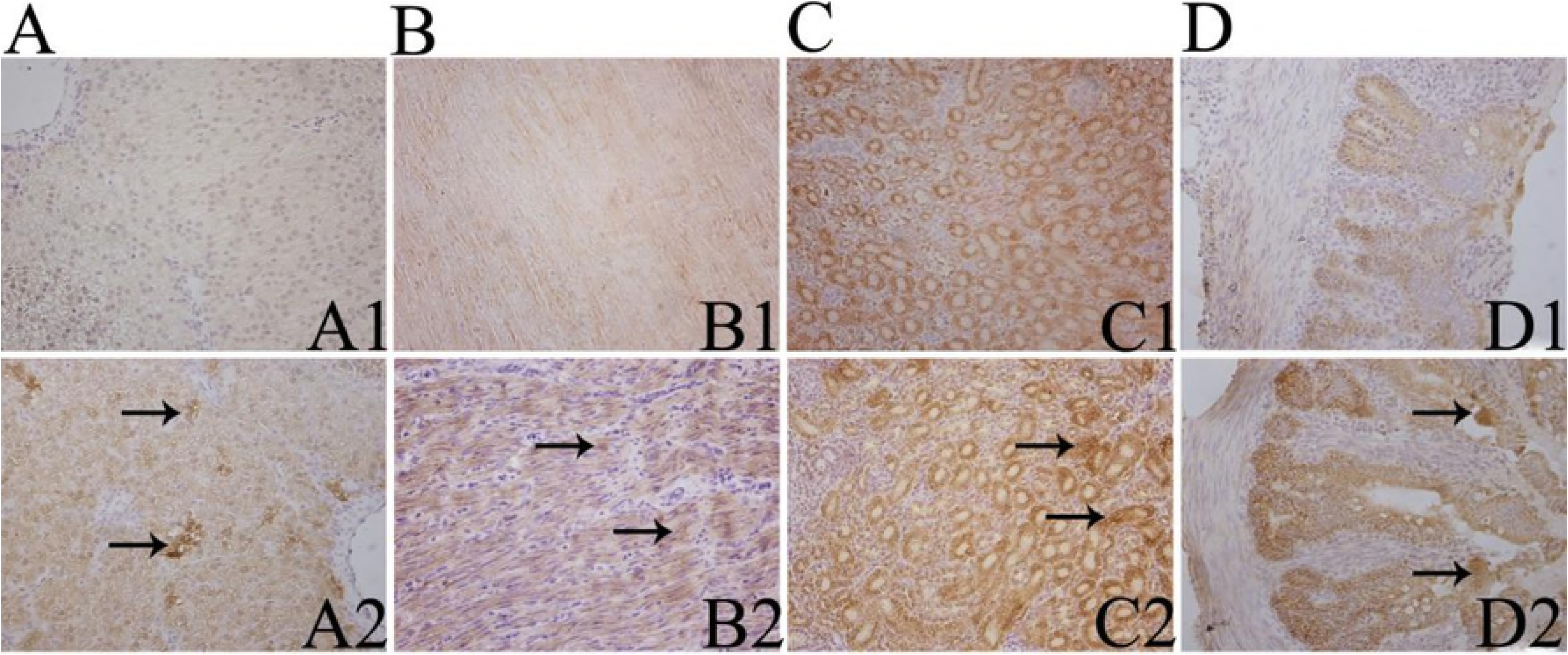
Histopathological observation of different tissues at different times PI (400X) A: Spleen. A1: control group at 4 h PI; A2: experimental group at 4 h PI; A3: control group at 48 h PI; A4: experimental group at 48 h PI. Compared with A1, the structure of the experimental group spleen was normal in A2. Compared with A3, lymphocyte numbers increased in A4 (Arrow). B: Brain. B1: control group at 8 h PI; B2: experimental group at 8 h PI; B3: control group at 48 h PI; B4: experimental group at 48 h PI. Compared with B1, the structure of the experimental group brain was normal in B2. Compared with B3, inflammatory cell infiltration is observed under the meningeal membranes in B4. C: Lung. C1: control group at 12 h PI; C2: experimental group at 12 h PI; C3: control group at 24 h PI; C4: experimental group at 24 h PI. Compared with C1, the pulmonary lobule showed serious compensatory dilation in C2. Compared with C3, the number of pulmonary capillaries decreased markedly in C4. D: Cecum. D1: control group at 24 h PI; D2: experimental group at 24 h PI; D3: control group at 216 h PI; D4: experimental group at 216 h PI. Compared with D1, the local intestinal mucosa showed necrosis in D2. Compared with D3, lymphocyte numbers increased in D4. E: Ileum, E1: control group at 48 h PI; E2: experimental group at 48 h PI; E3: control group at 72 h PI; E4: experimental group at 72 h PI. Compared with E1, the top of the villi epithelial cells showed necrosis in E2. Compared with E3, the inflammatory cells increased in E4. F: Duodenum. F1: control group at 72 h PI; F2: experimental group at 72 h PI; F3: control group at 216 h PI; F4: experimental group at 216 h PI. Compared with F1, the top of the villi epithelial cells showed necrosis in F2. Compared with F3, the villi showed erosion in F4.

**Fig 3.**
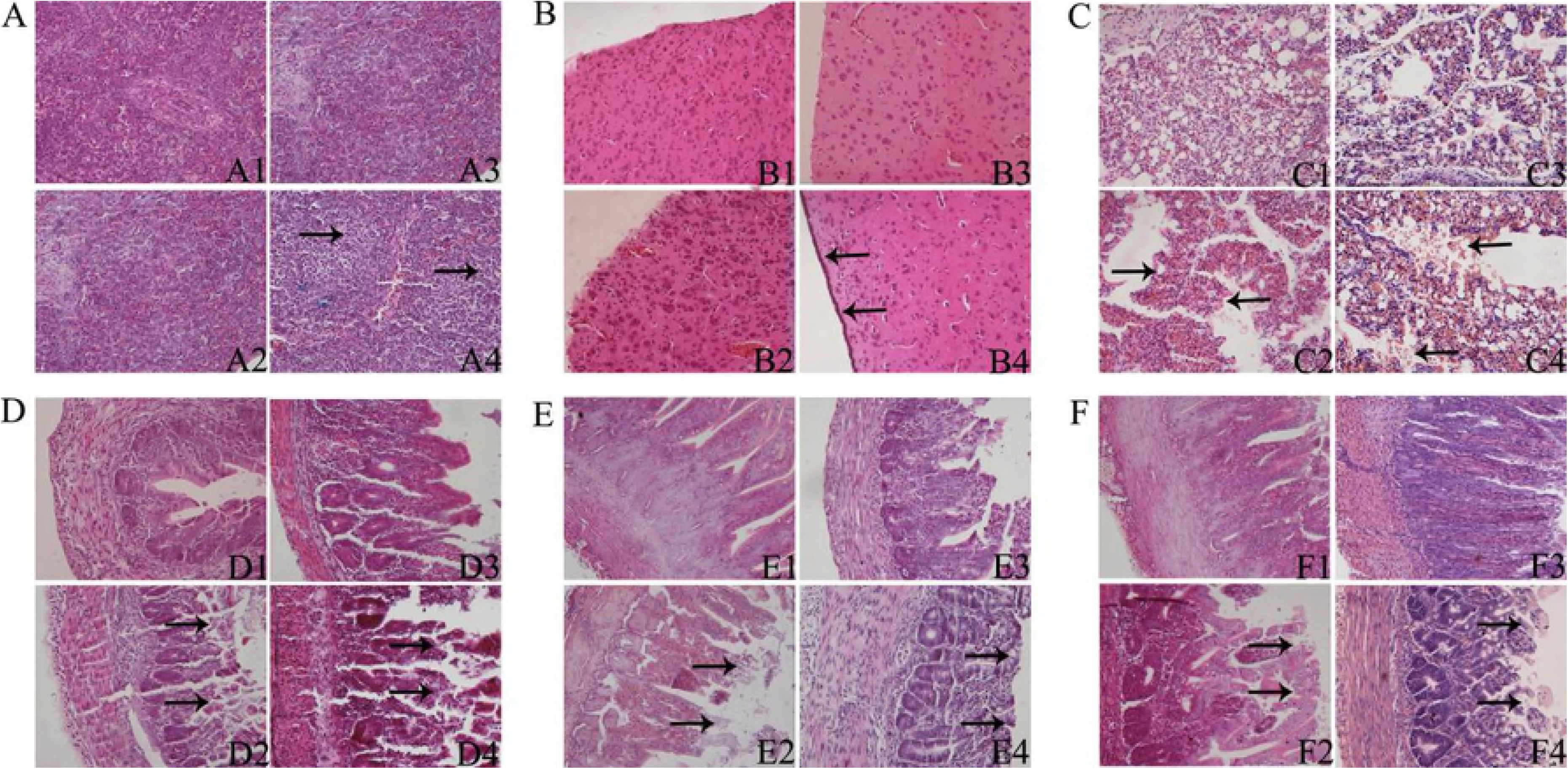
Histopathology of other tissues and organs after the Cherry Valley ducks were experimentally infected with duck-origin parvovirus. a: heart 400x; b: liver 400x; c: kidney 400x; d: pancreas 200x; e: thymus 200x; f: tongue 400x; g: proventriculus 200x; h: gizzard 400x; i: esophagus 200x; j: trachea 400x; k: jejunum 400x; l: rectum 400x. a1-ll are the control group; a2-l2 are the experimental group, and there was no obvious pathological damage.

**Fig 4.**
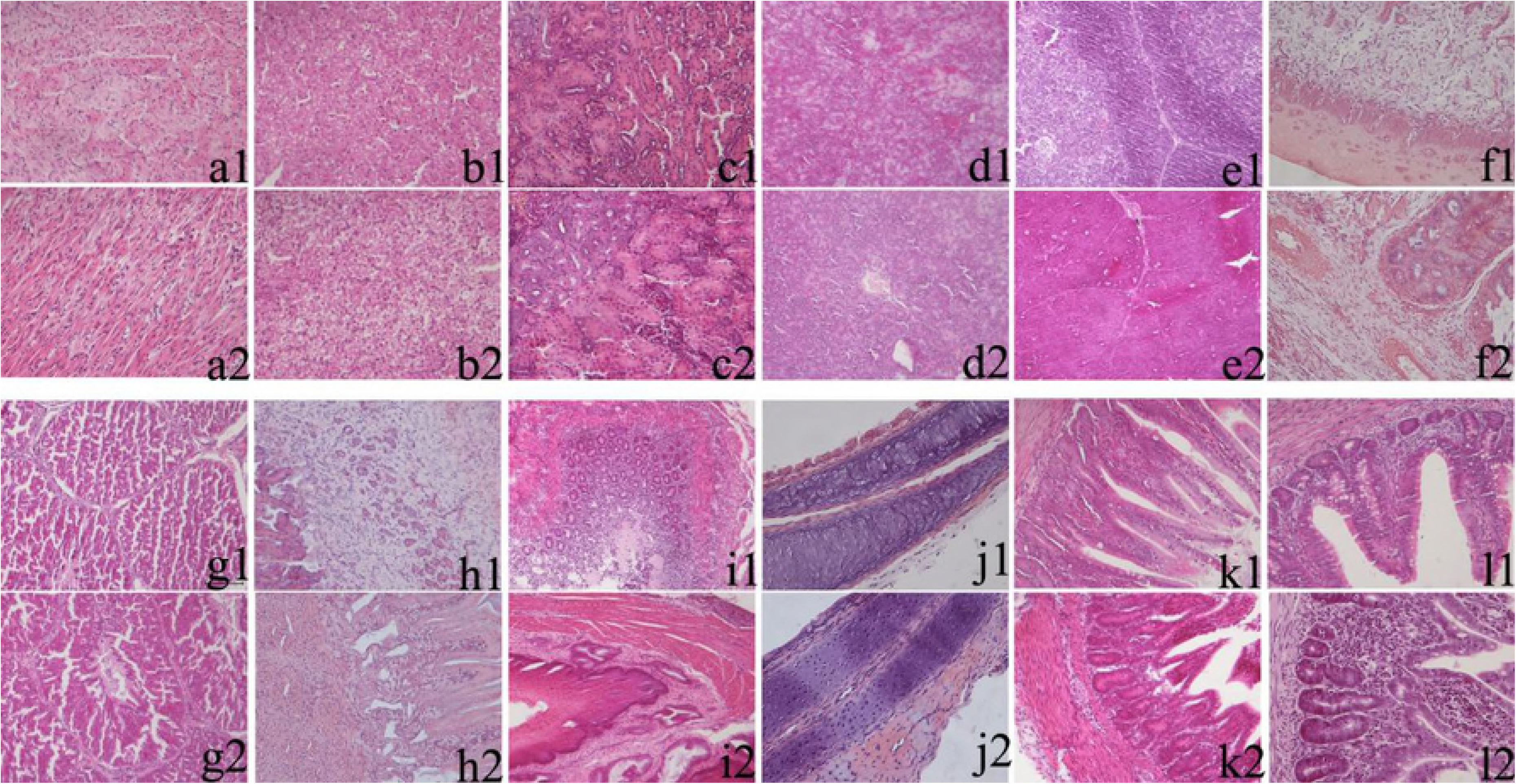
Viral antigen expression in different tissues and organs at different times PI (400X) A: Spleen. A1: control group at 4 h PI; A2: experimental group at 4 h PI. Positive expression is found in the white pulp and red pulp. B: Brain. B1: control group at 8 h PI; B2: experimental group at 8 h PI. Positive expression is found in brain cells. C:Lung. C1: control group at 12 h PI; C2: experimental group at 12 h PI. Positive expression is found in the epithelium of the lung atrium. D: Cecum. D1: control group at 24 h PI; D2: experimental group at 24 h PI. Positive expression is found in the mucosal epithelium. E: Ileum. E1: control group at 48 h PI; E2: experimental group at 48 h PI. Positive expression is found in the villus and intestinal crypt. F: Duodenum. F1: control group at 72 h PI; f2: experimental group at 72 h PI. Positive expression is found in the villus and intestinal crypt.

**Fig 5.**
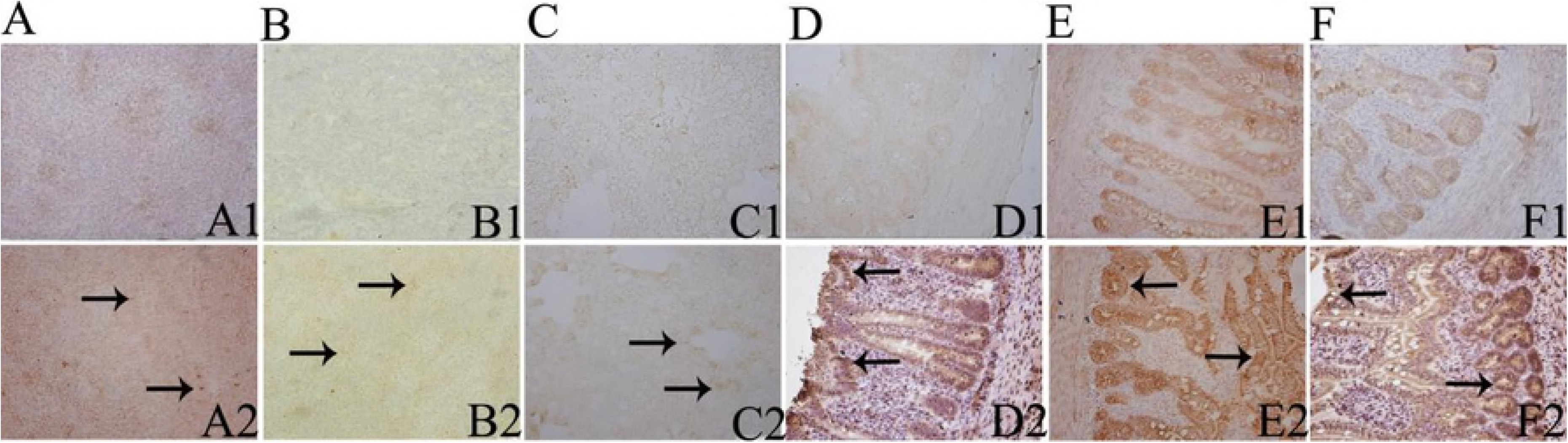
IHC of the liver, heart, kidney and jejunum after the Cherry Valley ducks were experimentally infected with duck-origin parvovirus (400X) A: Liver. A1: control group at 48 h PI; A2: experimental group at 48 h PI. Positive expression is found in the hepatic sinus and hepatocytes. B: heart. B1: control group at 48 h PI; B2: experimental group at 48 h PI. Positive expression is found in the myocardium. C: Kidney. B1: control group at 48 h PI; B2: experimental group at 48 h PI. Positive expression is found in the renal tubular epithelial cells. D: jejunum. D1: control group at 48 h PI; D2: experimental group at 48 h PI. Positive expression is found in the villus and intestinal crypt.

## 4. Discussion

Duck-origin parvovirus is a newly discovered parvovirus from recent years; Cherry Valley ducks infected with this virus display growth retardation, atrophy of the upper and lower beak and tongue swelling. According to reports, the study of duck-origin parvovirus has mainly focused on its isolation [4, 6-9], establishment of detection methods [24-25], and homology analysis [4, 9, 26], but the distribution of the virus in the duck and the main target organs have not been reported. Therefore, a detailed investigation of duck-origin parvovirus was carried out. In the present study, we used the duck-origin parvovirus QH-L01, which was successfully isolated from a Cherry Valley duck with characteristic signs of BADS, to systematically study the dynamic distribution of the virus in ducks and the damage to target organs caused by artificial infection, as well as to provide a clinical diagnosis and scientific basis for comprehensive prevention of this disease.

In the current study, duck-origin parvovirus (QH-L01) was first detected at 0.5 h PI in the blood with 10^2.01^ copies/mL, then in the spleen at 1 h with 10^2.86^ copies/g. Previous studies have examined the regular distribution pattern of GPV in geese and Muscovy ducks inoculated with GPV by qualitative PCR [14, 27]. Yang [14] found that GPV was first detected at 4 h PI in the liver and other organs. Compared with Yang, we advanced 3 h earlier to find the starting target of the duck-origin parvovirus infection in tissues. We observed that the copy numbers of QH-L01 in the brain, lung, cecum, ileum and duodenum were significantly higher than those in other tissues and organs at the same time point, suggesting that the virus accumulates or replicates in these organs. It is worth noting that the virus was detected in the blood but at low levels, suggesting that the duck-origin parvovirus is transported in the bloodstream without proliferation. Interestingly and in contrast to GPV, high viral load was detected in the lung at 12 h PI, showed that the lung may be a new target. In addition, at 24 h PI, the tissues with the highest copy numbers of QH-L01 were the cecum and the ileum and duodenum, at 48 h and 72 h-216 h, respectively. The results of IHC showed positive expression in the spleen, brain, lung and intestinal tissues, and the positive expression was localized in the intercellular spaces and epithelial cells, showing that the virus entered the cells and proliferated. These results further confirmed our hypothesis that a pathogenic process does occur, and we conclude that duck-origin parvoviruses are transmitted from the primary infection site (probably the spleen, as shown in this study) to the brain and lung, resulting in proliferation and dissemination of the virus to the cecum, ileum, duodenum and other tissues through the blood.

HE staining showed that the brain, lung, cecum, ileum and duodenum appeared to be damaged at different time points PI (Fig 2). At present, previous studies [1, 5, 9-10] on Cherry Valley Duck pathology in duck-origin parvovirus are limited to natural cases, but natural cases have random, uncontrolled infections and secondary infections from multiple pathogens, leading to unrepresentative pathological changes. In this study, the pathological changes in the brain, kidney and spleen were similar to those observed in natural cases, but the lung, duodenum, ileum and cecum were more obvious in this experiment. qPCR and IHC indicated that the spleen, brain, lung, cecum, liver, ileum and duodenum had high viral load or high expression over time (Fig 1,Fig 3). It was reported that target organ virus accumulation and proliferation play a role in toxic damage to organs [28-29]. Therefore, the brain, lung, cecum, ileum and duodenum can be considered the target organs during infection of healthy Cherry Valley ducks with duck-origin parvovirus. However, it is worth noting that the liver, heart, jejunum and other tissues throughout infection had high viral load or high antigen expression, but no pathological damage occurred at the same point in time. We hypothesized that the viral injury to tissues and organs was caused by virus proliferation or immune responses targeting viral antigens in the tissues [30]. Therefore, we speculated that the tissue/organ injury was directly caused by immune responses rather than by virus proliferation and was indirectly aggravated by viral proliferation inducing immune responses; however, confirmation will require further investigation and additional data.

In addition, severe clinical symptoms and death were initiated 2 days after inoculation of experimental ducks, and the copy number of QH-L01 in the lung reached 10^8.28^ copies/g at 12 h PI; meanwhile, pathological diagnosis showed that lung injury was enhanced at 24 h PI with significant lung housing atrophy and pulmonary capillary damage. Therefore, we conclude that the initial rapid proliferation of duck-origin parvovirus in the lung could damage the structure and function of lung tissue, leading to dyspnea and ultimately death. Alternatively, qPCR and IHC showed high-level viral loads in the intestine, and histological examination revealed necrosis and erosion of the villus tips in the intestine. The high viral load of QH-L01 indeed caused destruction of intestinal morphology and function, which resulted in diarrhea and nutritional absorption disorder in the infected Cherry Valley ducks. The autopsies revealed systemic hemorrhagic symptoms in the infected Cherry Valley ducks, and qPCR suggested that the blood contains the virus, which indicates that viremia occurred in Cherry Valley ducks infected with QH-L01. Over time, the copy number of QH-L01 decreased, and the lung, intestine, kidney and brain were gradually restored at 216 h PI. Histological examination of the spleen showed relative lymphocytosis at 48 h PI; thus, QH-L01 may stimulate the immune system through a feedback mechanism[31], thereby protecting Cherry Valley ducks from viral injury. Consequently, tolerance to infection at 72 h-144 h PI would lead to the survival of Cherry Valley ducks. Another interesting finding was the increase in intestinal lymphocytes before the splenic lymphocytes, which suggests that intestinal immunity was activated before splenic immunity; however, the reason for this result is unclear.

In summary, we demonstrated successful strategies for studying duck-origin parvovirus infection in Cherry Valley ducks by qPCR, HE staining and IHC. The results led us to conclude that duck-origin parvovirus can be transmitted from the spleen to the brain and lung, resulting in proliferation and dissemination of the virus to the cecum, ileum, duodenum and other tissues through the blood. Further, the brain, lung, cecum, ileum and duodenum, may represent the main target tissues/organs for the duck-origin parvovirus. The findings are very important for the development of targeted drugs that can rapidly provide strategic control of this virus.

## Author Contributions

B.C. carried out most of the experiments and wrote the manuscript. Z.C. and Q.L. critically revised the manuscript and the experimental design. J.X., C.H., W.L. and A.C helped with the experiments. All of the authors read and approved the final version of the manuscript.

